# Dock1 acts cell-autonomously in Schwann cells to regulate the development, maintenance, and repair of peripheral myelin

**DOI:** 10.1101/2023.10.26.564271

**Authors:** Ryan A. Doan, Kelly R. Monk

## Abstract

Schwann cells, the myelinating glia of the peripheral nervous system (PNS), are critical for myelin development, maintenance, and repair. Rac1 is a known regulator of radial sorting, a key step in developmental myelination, and we previously showed in zebrafish that loss of Dock1, a Rac1-specific guanine nucleotide exchange factor, results in delayed peripheral myelination in development. We demonstrate here that Dock1 is necessary for myelin maintenance and remyelination after injury in adult zebrafish. Furthermore, it performs an evolutionary conserved role in mice, acting cell-autonomously in Schwann cells to regulate peripheral myelin development, maintenance, and repair. Additionally, manipulating Rac1 levels in larval zebrafish reveals that *dock1* mutants are sensitized to inhibition of Rac1, suggesting an interaction between the two proteins during PNS development. We propose that the interplay between Dock1 and Rac1 signaling in Schwann cells is required to establish, maintain, and facilitate repair and remyelination within the peripheral nervous system.

## Introduction

Myelin, the lipid-rich multi-lamellar sheath that surrounds and insulates axons, plays a critical role in the vertebrate nervous system, enabling rapid transmission of nerve impulses (Jessen and Mirsky, 2005). In the peripheral nervous system (PNS), myelin is synthesized by Schwann cells (SCs), with each mature SC myelinating a single axonal segment (Monk et al., 2015). Derived from the neural crest, SCs progress through developmental stages delineated by the expression of specific genes and marked by significant morphological transformations (Ackerman and Monk, 2016). SC precursors (SCPs) undertake extensive longitudinal migration along pathfinding peripheral axons and subsequently differentiate into immature SCs, which engage in a specialized function known as radial sorting. During this process, an immature SC projects extensions into a bundle of axons and selectively identifies an individual axon to myelinate (Feltri et al., 2016). Following radial sorting, immature SCs that selected larger caliber axons enter a pro-myelinating state, enveloping and myelinating the chosen axon segment. Smaller caliber axons that don’t become myelinated associate with Remak SCs and form clusters of unmyelinated axons known as Remak bundles (Harty and Monk, 2017; Herbert and Monk, 2017). Proper regulation of SC homeostasis is required beyond development, where it is necessary to maintain myelin and function in repair and remyelination in the case of injury and disease (Bremer et al., 2011; Jessen and Mirsky, 2016; Jessen and Mirsky, 2019). While a large body of work has shed light on the multi-faceted functions SCs play throughout life (Taveggia and Feltri, 2022), a complete understanding of the signaling involved at each stage remains incompletely defined and represents a critical area for further exploration.

Work from our lab previously showed that Dock1, an evolutionarily conserved guanine nucleotide exchange factor (GEF), is required for timely radial sorting and developmental PNS myelination in zebrafish (Cunningham et al., 2018). Dock1 belongs to the 11-member family of Dock proteins, related in their ability to activate Rac1, fellow Rho-family member Cdc42, or a combination of both (Côté and Vuori, 2002). GEFs play a direct role in activating Rho-family GTPases in reaction to various extracellular signals and activity, enabling them to function as regulators of the cytoskeletal dynamics that underpin numerous cellular processes, ranging from migration, morphological changes, and phagocytosis (Côté and Vuori, 2002; Côté and Vuori, 2007; Hasegawa et al., 1996; Laurin et al., 2008; Rossman et al., 2005; Ruiz-Lafuente et al., 2015; Ziegenfuss et al., 2012). Additional work has begun to characterize the importance of several GEFs, including members of the Dock family, in regulating SC development and function (Miyamoto et al., 2016; Pasten et al., 2015; Yamauchi et al., 2008; Yamauchi et al., 2011). Dock1 specifically regulates the Rho-GTPase Rac1, an essential mediator of SC development, governing shape changes via regulation of the actin cytoskeleton (Kiyokawa et al., 1998; Nodari et al., 2008). During SC development, temporally varied levels of Rac1 sequentially control migration, commencement of radial sorting, and myelination. In a mouse model with SC-specific deletion of *Rac1*, SCs in developing sciatic nerves showed evidence of delayed radial sorting along with abnormal SC cytoplasmic extensions, ultimately resulting in severely delayed myelination (Benninger et al., 2007; Guo et al., 2012; Nodari et al., 2008). The function of Dock1 in the PNS is an emerging area of interest, and while it has been demonstrated to be required for proper PNS development in zebrafish, its roles in myelin maintenance, repair, and remyelination following injury remain unknown. Furthermore, *Dock1*’s high expression in the developing mouse PNS (Gerber et al., 2021) underscores its potential significance, yet its specific function in mammalian SCs has yet to be explored.

In this study, we employ zebrafish and mouse models to expand our knowledge of how Dock1 functions in the PNS. Our data reveal that Dock1 is instrumental for development but is dispensable for myelin maintenance into early adulthood in both zebrafish and mice. We show that aged animals in both species rely on Dock1 for the long-term maintenance of myelin integrity, with mature animals manifesting numerous aberrant myelin phenotypes. Moreover, we identify a critical function for Dock1 in the remyelination of axons after peripheral injury. Finally, manipulating Rac1 levels in development reveals an interaction with Dock1 that alters developmental myelination. Collectively, these findings illuminate Dock1’s complex and evolutionarily conserved role in SCs, where it regulates myelin development, homeostasis, and repair. Understanding the interplay between Dock1 and Rac1 may provide new insights into the pathways controlling myelin formation and maintenance, offering novel avenues for treating conditions that impact the peripheral nervous system.

## Results

### Dock1 functions in myelin maintenance in aged adult zebrafish

Our previously published work identified Dock1 as a regulator of developmental PNS myelination in zebrafish (Cunningham et al., 2018). Given that many genes required for myelin development are also necessary for myelin maintenance (Decker et al., 2006), we wanted to know if Dock1 played a role in myelin homeostasis into adulthood. To this end, we analyzed zebrafish maxillary barbels (ZMBs) using the previously described *dock1^stl145^* loss of function mutant zebrafish line. *stl145*, the allele designation of the *dock1* mutant our lab identified in a forward genetic screen, represents an early stop codon in the Rac1 binding domain of *dock1*. *dock1^stl145/+^* heterozygous mutants do not show any myelin phenotypes, while *dock1^stl145/stl145^*homozygous mutants exhibit delayed developmental myelination and have evidence of delayed radial sorting (Cunningham et al., 2018). Maxillary barbels are innervated sense organs found in fish, reptiles, and amphibians (Winokur, 1982). Zebrafish develop paired ZMBs at approximately one month of age (LeClair and Topczewski, 2009). They contain a variety of structures, including taste buds, goblet cells, and a population of pure sensory nerves branching from cranial nerve VII (LeClair and Topczewski, 2009; LeClair and Topczewski, 2010; Moore et al., 2012). We performed ultrastructural analyses of ZMBs from 4-month-old and 1-year-old wild-type (WT), *dock1^stl145/+^*heterozygous, and *dock1^stl145/stl145^* homozygous mutant animals by transmission electron microscopy (TEM). At four months, we observed no changes in either the number of myelinated axons or in g-ratios, nor did we note any obvious myelin defects in heterozygous *dock1^stl145/+^* or homozygous *dock1^stl145/stl145^* mutant ZMBs compared to WT *dock1^+/+^* controls (Fig. 1, A-C and Fig S1, A-D). At one year, however, we found that there was a significant increase in the percentage of abnormal myelinated axons profiles in homozygous mutants compared to WT and heterozygous controls (*dock1^+/+^* (WT) 4 mo. old = 1.88%, *dock1^stl145/stl145^* (Mutant), 4 mo. old = 2.22%, WT 1 yr. old. = 4.78%, *dock1* 1 yr. old = 8.56%, P = 0.0002; Fig. 1, D-I). These findings suggest that Dock1 is necessary for long-term myelin maintenance in zebrafish but is dispensable during early adulthood.

**Figure 1.**
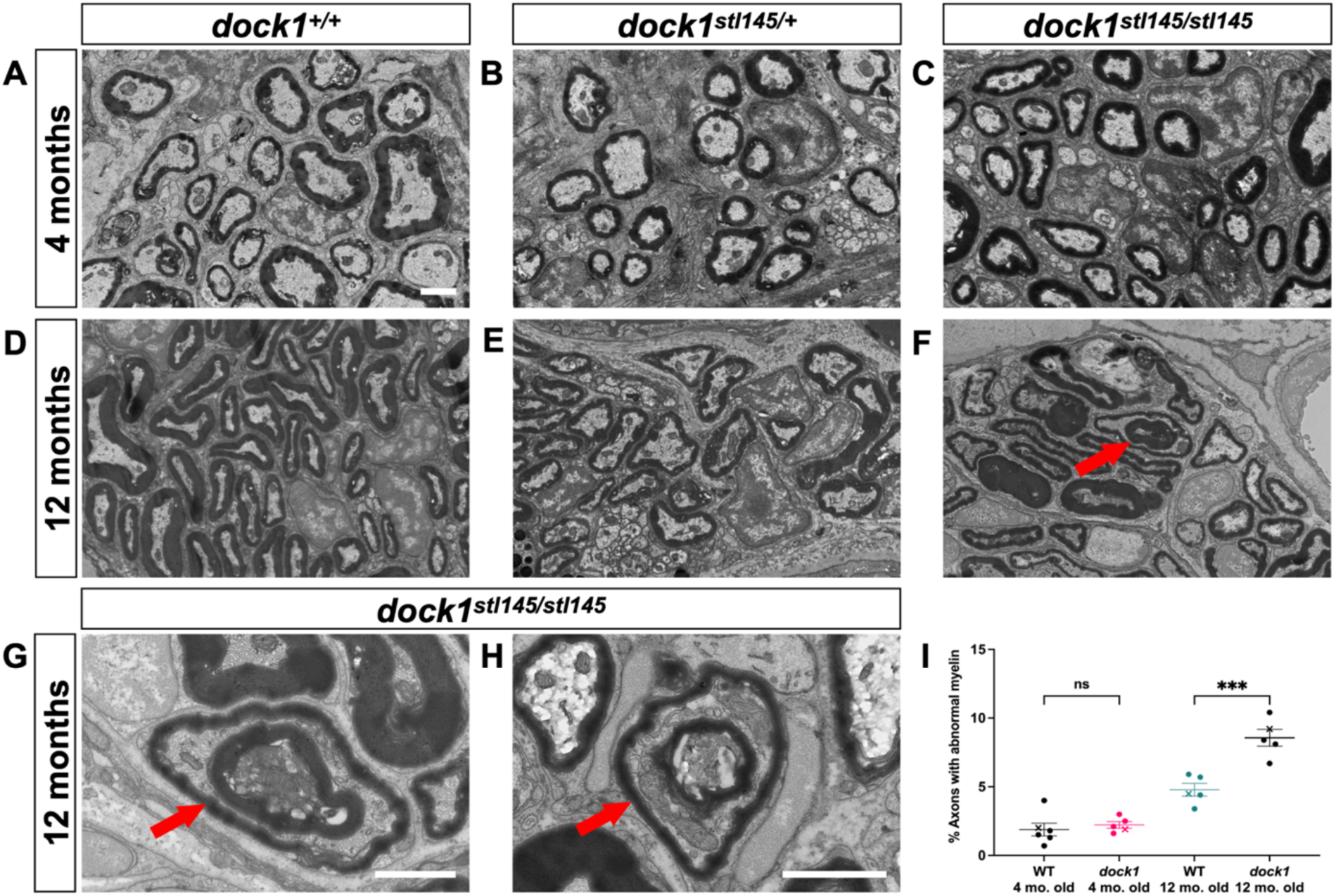
Age-dependent myelin maintenance defects are present in *dock1* mutants. **(A-C)** Transmission electron micrographs (TEM) of cross-sections of zebrafish maxillary barbels (ZMBs) from 4-month-old WT *dock1^+/+^*, heterozygous *dock1^stl145/+^*, and homozygous *dock1^stl145/stl145^* mutant zebrafish. **(D-F)** TEM micrographs of ZMBs from 12-month-old WT *dock1^+/+^*, heterozygous *dock1^stl145/+^*, and homozygous *dock1^stl145/stl145^*mutant zebrafish, with homozygous mutants exhibiting myelin outfoldings (red arrow). **(G, H)** Higher magnification TEM micrographs of ZMBs from 12-month-old homozygous *dock1^stl145/stl145^*mutants showing abnormally myelinated axons (red arrows), features rarely seen in WT *dock1^+/+^* or heterozygous *dock1^stl145/+^* mutants. **(I)** Quantification of the percent of axons with abnormal myelin profiles, observed by TEM, in WT *dock1^+/+^*vs homozygous *dock1^stl145/stl145^* mutants at 4-months and 12-months-old, *n = 10* (WT *dock1^+/+^* 4 mo. old), 10 (*dock1^stl145/stl145^*mut. 4 mo. old), 10 (WT *dock1^+/+^*12 mo. old), 10 (*dock1^stl145/stl145^* mut 12 mo. old). The “X” symbol in the graph denotes a data point corresponding to the representative image shown. **(A-H)** Scale bar = 1 µm. **(I)** Two-way ANOVA with Tukey’s multiple comparisons test. ***, P < 0.0005; ns, not significant.

### Remyelination following injury is significantly impaired in *dock1* mutant zebrafish

To further our understanding of the role of Dock1 in the developed PNS, we examined its role in remyelination following injury in zebrafish. The ZMB can regrow after amputation, and axons with nerves of the regenerating ZMB are remyelinated as the appendage regrows, with myelin reaching around 85% of its original thickness 4-weeks following transection (Moore et al., 2012). We, therefore, removed the left ZMB from 3-month-old animals (a timepoint when myelin maintenance defects are not yet apparent) via cut and allowed recovery for 4 weeks. The right uncut ZMB served as an internal uninjured control for each animal. After 4 weeks, ZMBs from both sides were removed, processed for TEM, and the nerves were examined. The regenerated barbels of the WT, heterozygous (data not shown), and homozygous *dock1* mutants were similar in appearance and had both regrown to ∼90% of their original length (Fig. S2, A-D), suggesting that Dock1 is not required for gross ZMB regeneration. However, we observed a profound loss in the number of myelinated axons in the regenerated ZMBs of *dock1* mutants compared to WT (*dock1^+/+^* control = 19.85, *dock1^+/+^* regenerated = 16.87, *dock1^stl145/stl145^* control = 16.82, *dock1^stl145/stl145^* regenerated = 4.80, P = 0.0013; Fig. 2, A-E). Moreover, the myelin that was present in the mutants was much thinner than in controls as analyzed by g-ratio (Fig. 2, F). The number of SC nuclei and total axons were quantified by examining TEM micrographs, and we observed no differences in these parameters between regenerated WT and *dock1* mutant ZMBs (Fig. S2, E and F). These results suggest a crucial function for Dock1 in regulating remyelination of the PNS following nerve injury in zebrafish.

**Figure 2.**
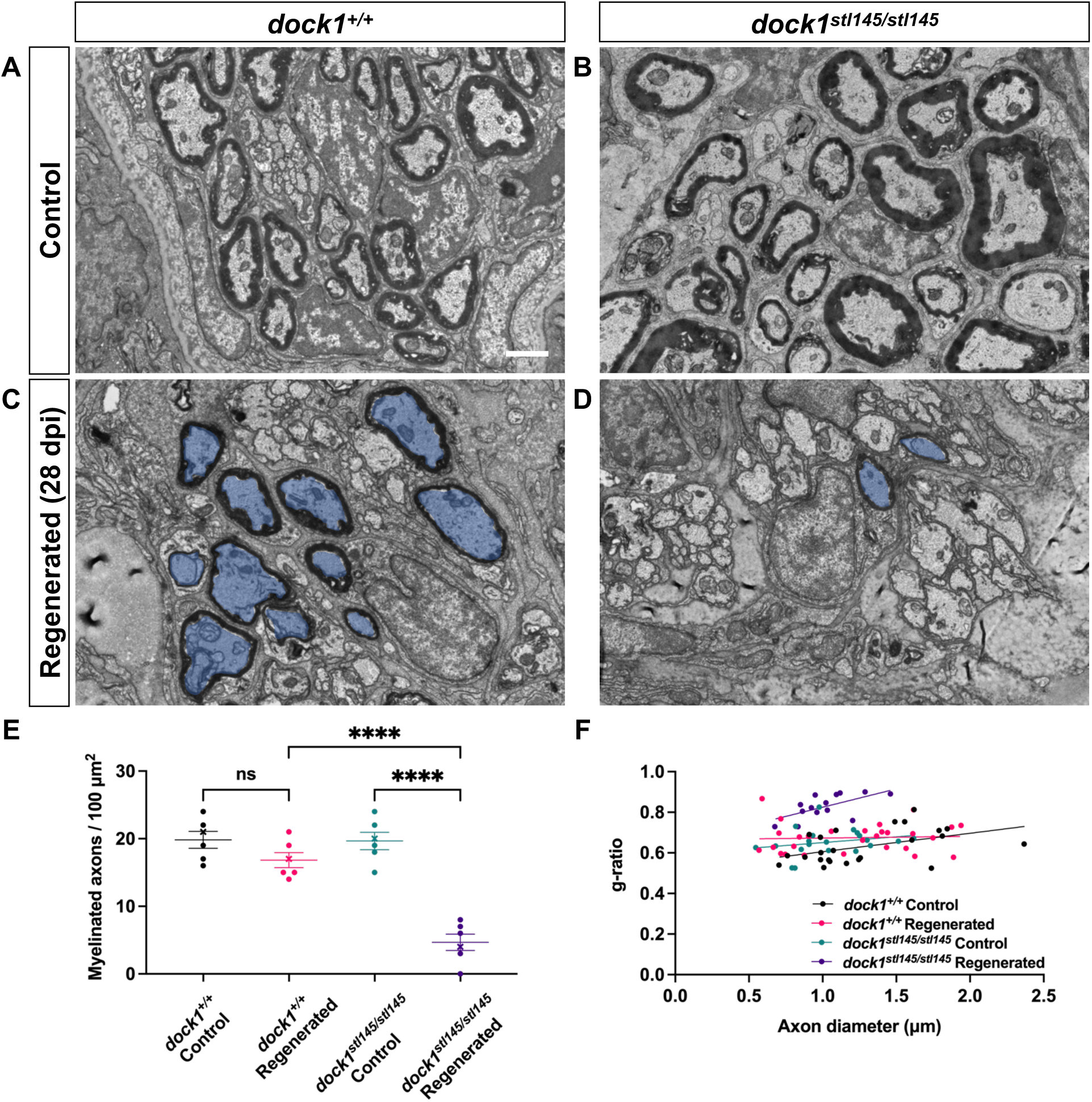
Remyelination following nerve injury is significantly reduced in *dock1* mutants. **(A-B)** TEM micrographs of control ZMBs from 4-month-old WT *dock1^+/+^* and mutant *dock1^stl145/stl145^* zebrafish. **(C, D)** TEM micrographs of WT *dock1^+/+^* and mutant *dock1^stl145/stl145^* zebrafish showing regeneration and remyelination after transection, with remyelinated axons pseudocolored in blue. **(E)** Quantification of the number of myelinated axons in the ZMBs per 100 µm^2^, *n = 6* (WT *dock1^+/+^* control), 6 (WT *dock1^+/+^* regenerated), 6 (*dock1^stl145/stl145^* mutant control), 6 (*dock1^stl145/stl145^* mutant regenerated). **(F)** Quantification of the g-ratio as it relates to axon caliber of the remyelinated axons in the regenerated ZMBs, 28 days after transection, *n = 6* (WT *dock1^+/+^*control), 6 (WT *dock1^+/+^* regenerated), 6 (*dock1^stl145/stl145^* mutant control), 6 (*dock1^stl145/stl145^* mutant regenerated). **(A-D)** Scale bar = 1 µm **(E)** Two-way ANOVA with Tukey’s multiple comparisons test. ****, P < 0.0001; ns, not significant.

### Dock1 functions cell-autonomously in Schwann cells to regulate myelination

Our work in global zebrafish *dock1* mutants demonstrates a vital role for Dock1 in regulating PNS myelin, from development to maintenance and repair. We hypothesized that loss of Dock1 function specifically in SCs is responsible for the phenotypes we observe in zebrafish for three reasons: 1) Dock1 is highly expressed in developing SCs (Gerber et al., 2021); 2) The known link between Dock1 and Rac1 signaling (Côté and Vuori, 2007); 3) The importance of Rac1 signaling in SCs (Abu-Thuraia et al., 2015). To test this theory and simultaneously determine if the function of Dock1 is evolutionarily conserved in mammals, we generated SC-specific *Dock1* conditional knockout (cKO) mice by crossing validated *Dock1^fl/fl^* mice (Laurin et al., 2008) with the well-characterized *Dhh^Cre^* mouse line (Jaegle et al., 2003) to drive recombination in SCPs at approximately embryonic day (E)12.5. Western blotting revealed a ∼70% reduction in Dock1 protein levels in sciatic nerve of *Dhh^(+)^;Dock1^fl/fl^* mice compared to littermate controls (Fig. 3 A; Fig. S3, A-B). We first examined the sciatic nerves of animals on postnatal day (P)3, when radial sorting is actively underway (Ackerman and Monk, 2016). Ultrastructural analyses by TEM revealed that *Dock1* cKO animals had significantly thinner myelin than their littermate controls (Fig. 3, B-D). To determine if this was due to a broader developmental defect in the SCs or the nerve itself, we examined TEM images and quantified the number of SC nuclei, unmyelinated axons, and myelinated axons. We found no significant differences between groups (Fig. S3, C-E). Upon closer examination of higher magnification TEM micrographs, we noticed that the SCs in the mutant animals exhibited additional defects. These included elaborate cytoplasmic protrusions extending from mutant SCs (Fig. 3 E), and evidence of basal lamina trails in regions devoid of SC cytoplasm (Fig. 3 F), suggesting that unstable SC process extensions had been made and retracted (Benninger et al., 2007; Nodari et al., 2007). To determine if these defects persisted throughout development, we examined animals at P28, when radial sorting is complete, and most myelin is established (Ackerman et al., 2018). Interestingly, at P28, *Dock1* cKO animals appear indistinguishable from WT controls (Fig. S3, F and G). There was no difference in myelinated axon number of (Fig. S3, H), and the increased g-ratio observed in the mutants at P3 had resolved (Fig. S3, I). These findings parallel what we observed in zebrafish and reveal that Dock1 is an evolutionarily conserved regulator of developmental myelination due to its function in SCs.

**Figure 3.**
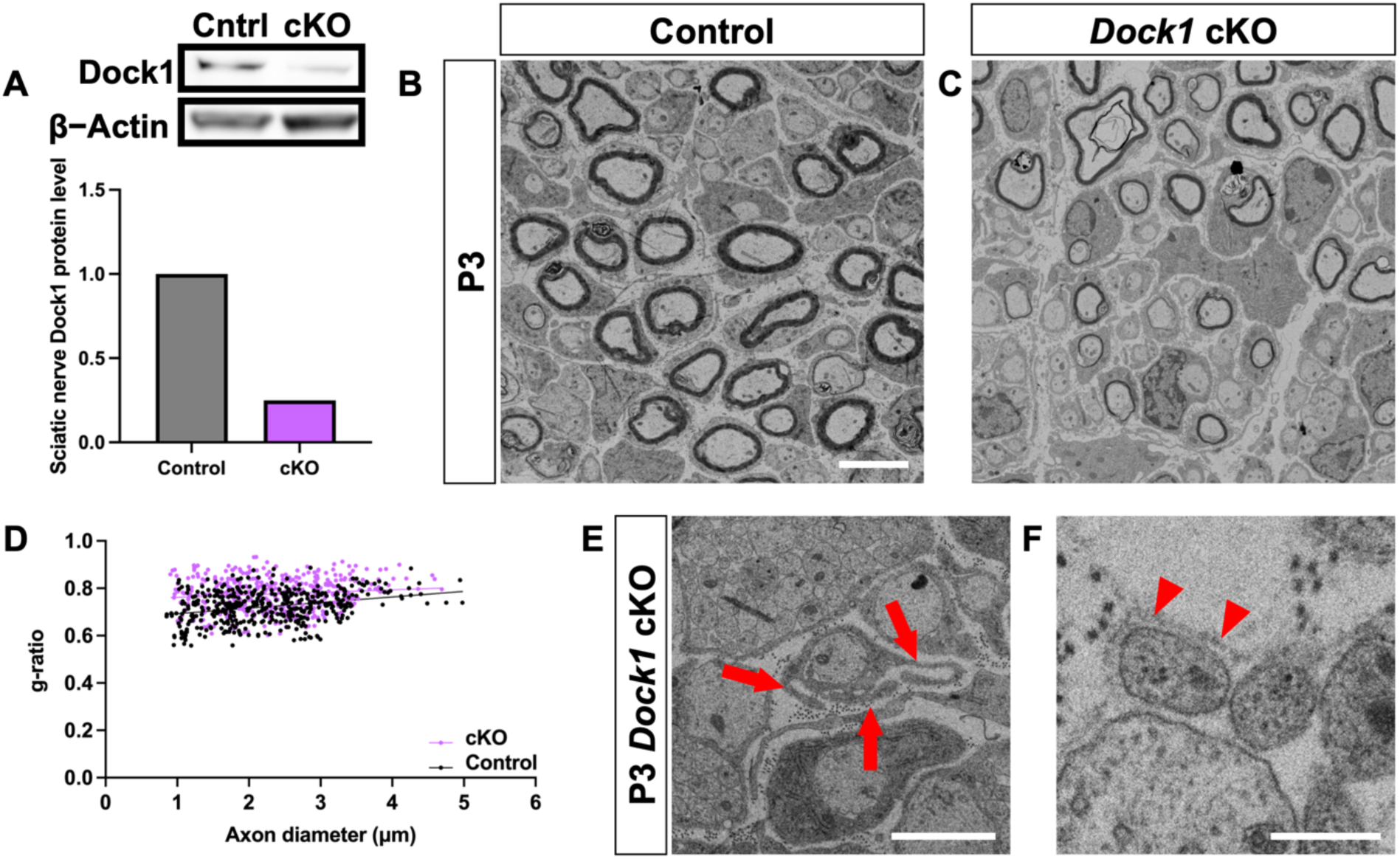
Schwann cell specific Dock1 mutants present with multiple defects in peripheral nerves. **(A)** Western blot showing sciatic nerve Dock1 and β-actin protein levels from control and *Dock1* cKO animals and quantification of normalized protein levels. **(B-C)** TEM micrographs of sciatic nerves from *Dhh^Cre+^;Dock1^+/+^*control and littermate *Dhh^Cre+^;Dock1^fl/fl^* cKO mice at postnatal day (P)3. **(D)** Quantification of the g-ratio as it relates to axon caliber, *n =* 6 mice, 4 images per nerve (wildtype), 4 mice, 4 images per nerve (cKO). **(E)** *Dock1* cKO mutant SCs display abnormal cytoplasmic protrusions that extend in multiple directions (red arrows). **(F)** Trails of basal lamina found in *Dock1* cKO mutants are observed in regions devoid of SC cytoplasm (red arrowheads). **(B, C)** Scale bar = 4 µm, **(E, F)** Scale bar = 1 µm.

### *Dock1* SC-specific knockout mice show age-associated myelin abnormalities

Proper maintenance of SCs in the developed PNS is essential for these cells to support the physiological health of the adult. When mature SC homeostasis is disrupted, it can present with various abnormalities, including muscle atrophy, decreased nerve conduction velocities, and sensory loss (Verdú et al., 2008). Several mutants with abnormal SC development are often accompanied by lifelong myelin defects (Bremer et al., 2011; Decker et al., 2006). In contrast, the delayed radial sorting and developmental hypomyelination seen in our *Dock1* mutants resolved as early as P28. Some mutants, such as *Gpr56/Adgrg1*, have a similar pattern of developmental SC defects that recover by early adulthood but show myelin abnormalities with age (Ackerman et al., 2018). To determine if this was the case for Dock1, we performed ultrastructural analyses of mouse sciatic nerves at 12 months. TEM revealed numerous myelin abnormalities in the aged 12-month-old *Dock1* cKO mutants compared to their younger P28 counterparts and age-matched littermates, including abnormal myelinated fibers and Remak defects (Fig. 4, A and B). We saw signs of degenerating myelin sheaths and accumulated axonal debris (Fig. 4, C-E), as well as regeneration clusters (Fig. 4, F) and myelin outfoldings (Fig. 4, G). Although control mice also showed some defects with age, these abnormalities were significantly more prevalent in mutants (Control 12 month = 4.02% of axons with abnormal myelin profiles, *Dock1* cKO 12 month = 12.41% of axons with abnormal myelin profiles, P = 0.0027; Fig. 4, H). These findings indicate that Dock1 is required in mouse SCs for long-term myelin maintenance and axonal health, and align with our observations in zebrafish, where myelin is normal in early adulthood, but defects arise with age.

**Figure 4.**
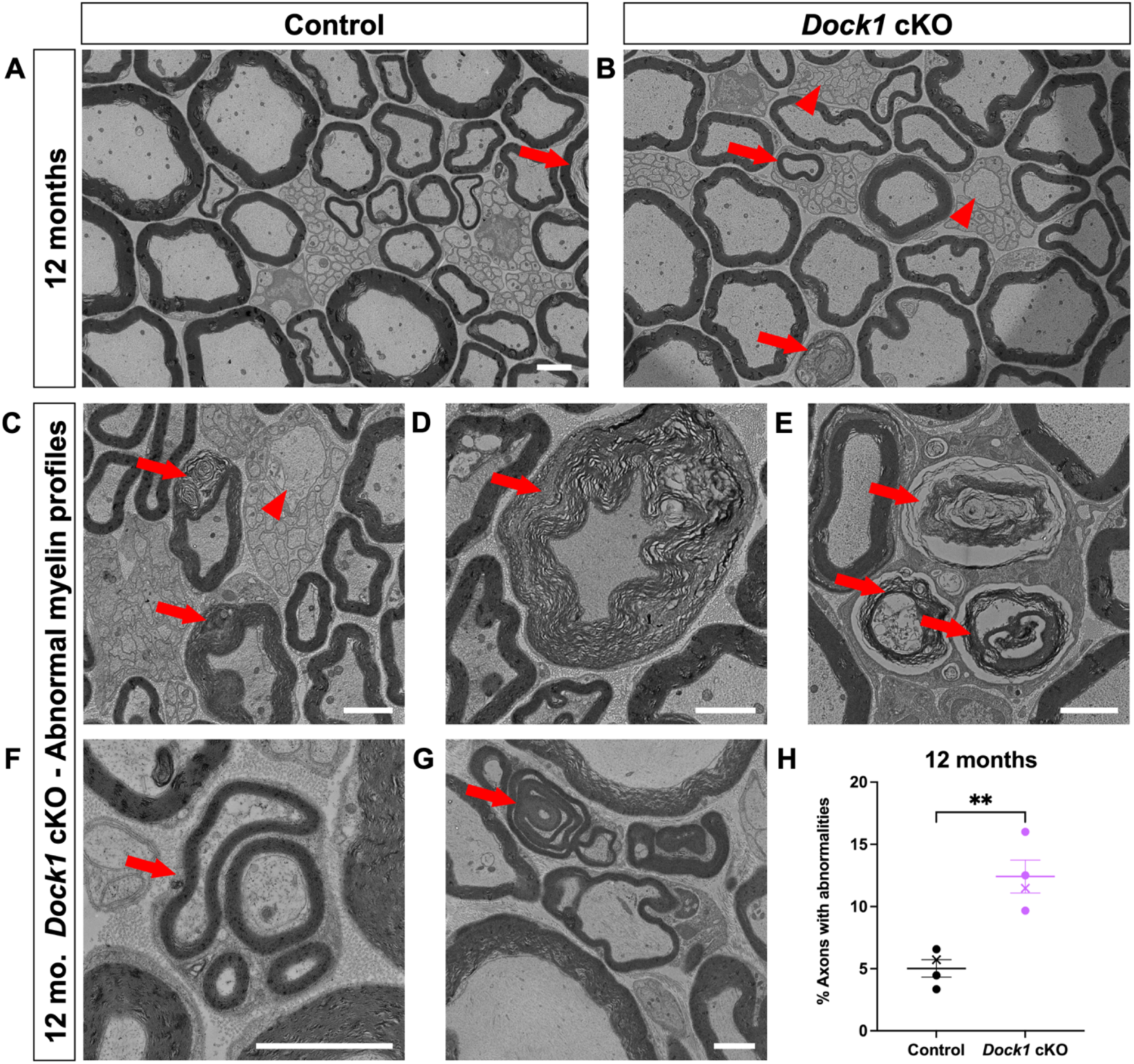
Myelin maintenance defects arise and accumulate with age in *dock1* cKO mice. **(A, B)** TEM micrographs of control and mutant nerves at 12 months with mutants containing aberrant myelin (red arrows) and Remak (red arrowheads) phenotypes. The following higher magnification TEM micrographs are from 12-month-old *Dock1* cKO nerves: **(C)** Degenerating axon (red arrow) and a large caliber (>1 μm) axon in a Remak bundle (red arrowhead) **(D)** Disorganized myelin sheath (red arrow) **(E)** Accumulations of myelin debris and degenerating sheaths (red arrows). **(F)** Regeneration clusters (red arrow) **(G)** Myelin outfoldings and abnormal wrapping (red arrow) **(H)** The percentage of axons with abnormalities, *n =* 3 mice, 4 images per nerve (control), 3 mice, 4 images per nerve (*Dock1* cKO). **(A-G)** Scale bar = 2 µm. **(H)** Unpaired t test with Welch’s correction. **, P < 0.005.

### Regeneration and remyelination are impaired in inducible *Dock1* SC-specific knockout mice

To help integrate the findings from our ZMB injury model and the cell-autonomous role of Dock1 in SCs, we examined its importance in mammalian remyelination. To assess this, we used the well-characterized mouse line *Plp^CreERT2^*(Leone et al., 2003) to generate an inducible conditional knock-out (icKO) mouse, allowing us to disrupt *Dock1* in mature SCs, leaving it functional during development. To assess repair after injury, we performed sciatic nerve transections, where the role of the SCs in regeneration and remyelination has been well described (Jessen and Mirsky, 2016). We transected the sciatic nerves of 3-month-old tamoxifen-injected *Plp^Cre+^;Dock1^fl/fl^* (icKO) mice and corn oil-injected *Plp^Cre+^;Dock1^fl/fl^*(control) mice, 4 weeks following the final tamoxifen injection, allowing sufficient time for recombination (Fig. S4, A) (Leone et al., 2003; Mogha et al., 2016). Following nerve transection, a bridge rapidly forms, and the nerve regenerates, making it difficult to see the injury site without resorting to immunostaining (Cattin and Lloyd, 2016; Dun and Parkinson, 2015). To ensure we examined nerves at the same distance from the cut site when we performed TEM, we quickly crushed the nerve with forceps coated in activated charcoal to mark the cut site before transection. We examined and analyzed the nerves at 14- and 25-days post-injury (dpi), time points that allow us to assess the clearance of debris associated with degenerating axons and also remyelination, respectively (Wang et al., 2023). Western blotting revealed a ∼40% reduction in Dock1 protein levels in the sciatic nerve of our tamoxifen-injected *Plp^(+)^;Dock1^fl/fl^* mice compared to corn oil injected controls (Fig. S4, B). When we examined the uninjured nerves of the 4-month-old icKO mice, we saw that the myelin abnormalities observed at 12 months in the cKO mice had yet to arise (Fig. 5, A and B). At 14 dpi, control mice showed the hallmarks associated with debris clearance, such as macrophages with internalized debris, which create an environment more conducive for axon regeneration. In contrast, SCs in the *Dock1* icKO mice were disorganized and had foamy macrophages (Fig. 5, C and D). By 25 dpi, control animals showed axons of various calibers that had begun to be remyelinate (Fig. 5, E). In contrast, icKO nerves had fewer remyelinated axons, and those present had higher g-ratios, despite having similar numbers of >1µm regenerated axons large enough to potentially be remyelinated compared to control mice (Control = 6.69, icKO = 2.21, P = 0.0021; Fig. 5, F-I). Our findings in this sciatic nerve transection model complement what we observed in ZMB regeneration and further support the conclusion that Dock1 is critical for SCs to regulate remyelination following peripheral nerve injury.

**Figure 5.**
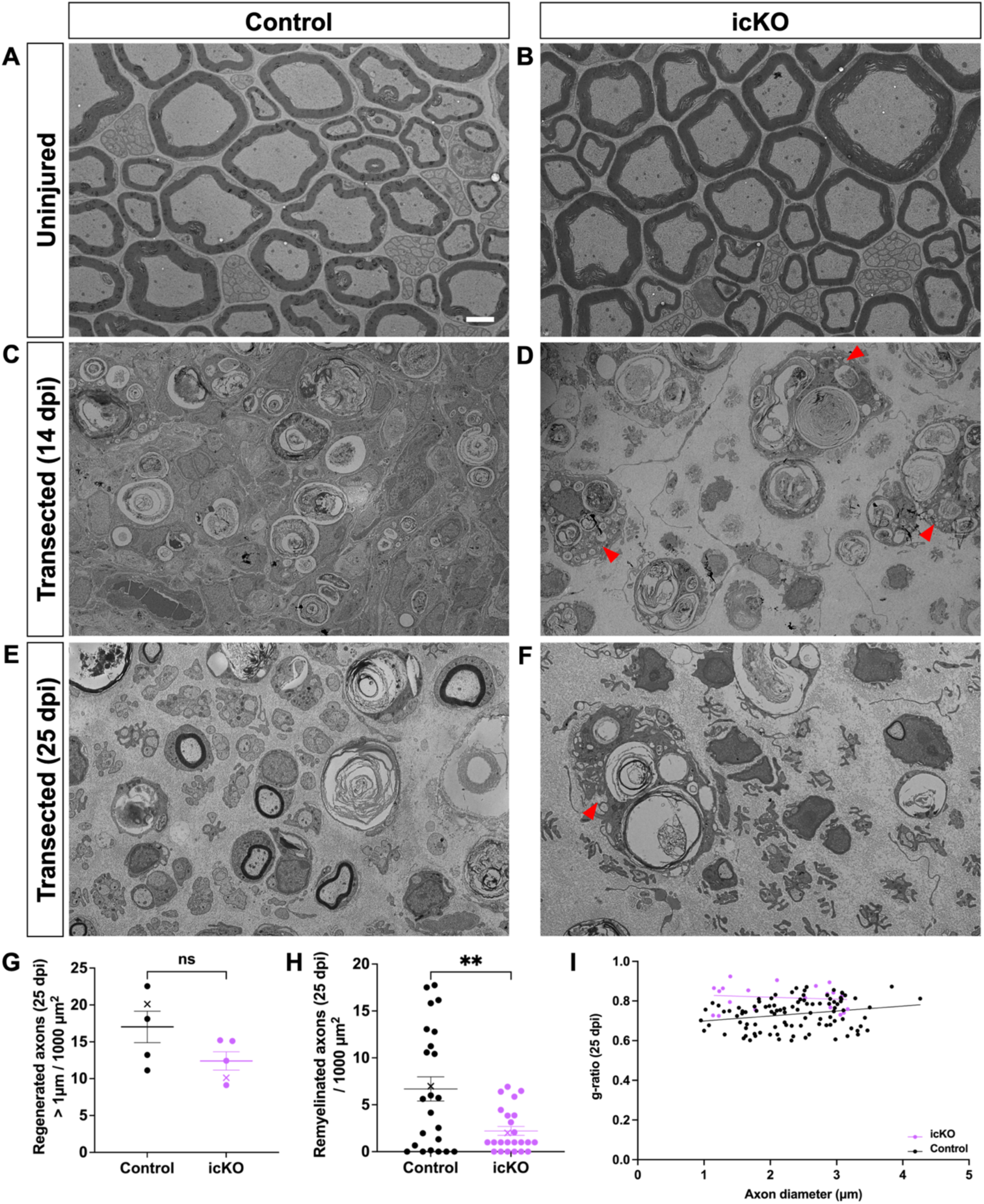
Remyelination is delayed following sciatic nerve transection in *Dock1* icKO mice. **(A-F)** TEM micrographs of sciatic nerves from control-injected and tamoxifen-injected *Plp^Cre+^;Dock1^fl/fl^* mice before injury, 14 days post-transection, and 25 days post-transection. **(G)** Quantification of the number of regenerated axons > 1 µm per 1000 µm^2^ **(H)**, the number of remyelinated axons per 1000 µm^2^, and **(I)** the g-ratio as it relates to axon caliber between control- and tamoxifen-injected mice, *n =* 6 mice, 4 images per nerve (control), 6 mice, 4 images per nerve (tamoxifen). **(A-F)** Scale bar = 2 µm. **(G, H)** Unpaired t test with Welch’s correction. **, P < 0.005; ns, not significant.

### Rac1 inhibition enhances myelin defects in *dock1* mutants

Rac1 is essential in SCs for radial sorting and myelination (Benninger et al., 2007; Nodari et al., 2007). Given Dock1’s GEF activity for Rac1, we wanted to know whether manipulating Rac1 levels would alter myelination in *dock1^stl145/+^* heterozygotes, which are typically indistinguishable from WT, or enhance the *dock1^stl145/stl145^* mutant hypomyelination phenotype. We performed a pharmacological sensitization study using the Rac1 inhibitor EHT1864 (Onesto et al., 2008) and used whole mount *in situ* hybridization (WISH) for *myelin basic protein* (*mbp*) to assess *mbp* expression in the developing posterior lateral line (PLLn). The PLLn is a major peripheral sensory nerve that runs the length of the zebrafish and begins myelinating around 3 days post-fertilization (dpf) (Sarrazin et al., 2010). We previously showed that *dock1^stl145/stl145^* mutants have a slight reduction in *mbp* expression in the PLLn at 5 dpf compared to WT (Cunningham et al., 2018). Consistent with our prior work, at 4 dpf, *dock1^stl145/stl145^*mutants also have slightly reduced *mbp* expression in the PLLn compared to WT, while *dock1^stl145/+^* heterozygotes are indistinguishable from WT (Fig. 6, A-C). A dose-response study was done by administering EHT1864 from 2-4 dpf, and we found that at 5 µM, there was no effect on the overall health of WT zebrafish, whereas higher doses resulted in toxicity. Treating zebrafish from 2-4 dpf allows us to target SCs during the onset of radial sorting and the initiation of myelination. Upon examining the PLLn at 4 dpf, we saw that *mbp* expression in PLLns from WT zebrafish were unaffected by the 5 µM dose of EHT1864 (Fig. 6, D). *dock1^stl145/+^* heterozygotes, however, showed a reduction in *mbp* expression compared to the treated WT and untreated controls (Fig. 6, E). The same was true for *dock1^stl145/stl145^* mutants but to an even greater extent, with some segments of the PLLn completely devoid of *mbp* expression (Fig. 6, F). When we quantified our observations, we found that there were no significant differences in *mbp* expression between WT and *dock1^stl145/+^* heterozygotes in our DMSO-treated controls (Fig. 6, G), however, there was a significant correlation between the phenotypes observed and the genotypes in the EHT1864-treated zebrafish (Fig. 6, H). Next, we examined myelin ultrastructure by performing TEM. At 4 dpf, *dock1^stl145/stl145^* mutants had a slight reduction in the number of myelinated axons at baseline compared to WT and *dock1^stl145/+^* heterozygotes (Fig. 6, I-K). By TEM, EHT1864-treated WT zebrafish didn’t have a discernable change in the number of myelinated axons compared to the untreated controls (Fig. 6, L). In contrast, there was a reduction in the number of myelinated axons in the *dock1^stl145/+^* heterozygous and *dock1^stl145/stl145^* mutants following drug treatment (Fig. 6, M and N). When we quantified the total number of axons, we saw no difference between any of the genotypes before or after 5 µM EHT1864 treatment (Fig. 6, O); however, when we analyzed the number of myelinated axons, we saw that *dock1^stl145/+^* heterozygous and *dock1^stl145/stl145^* mutants are sensitized to Rac1 inhibition (Fig. 6, P), providing support for the concept that even modest disruption to Dock1-Rac1 signaling can result in dysregulated myelination in the developing PNS.

**Figure 6.**
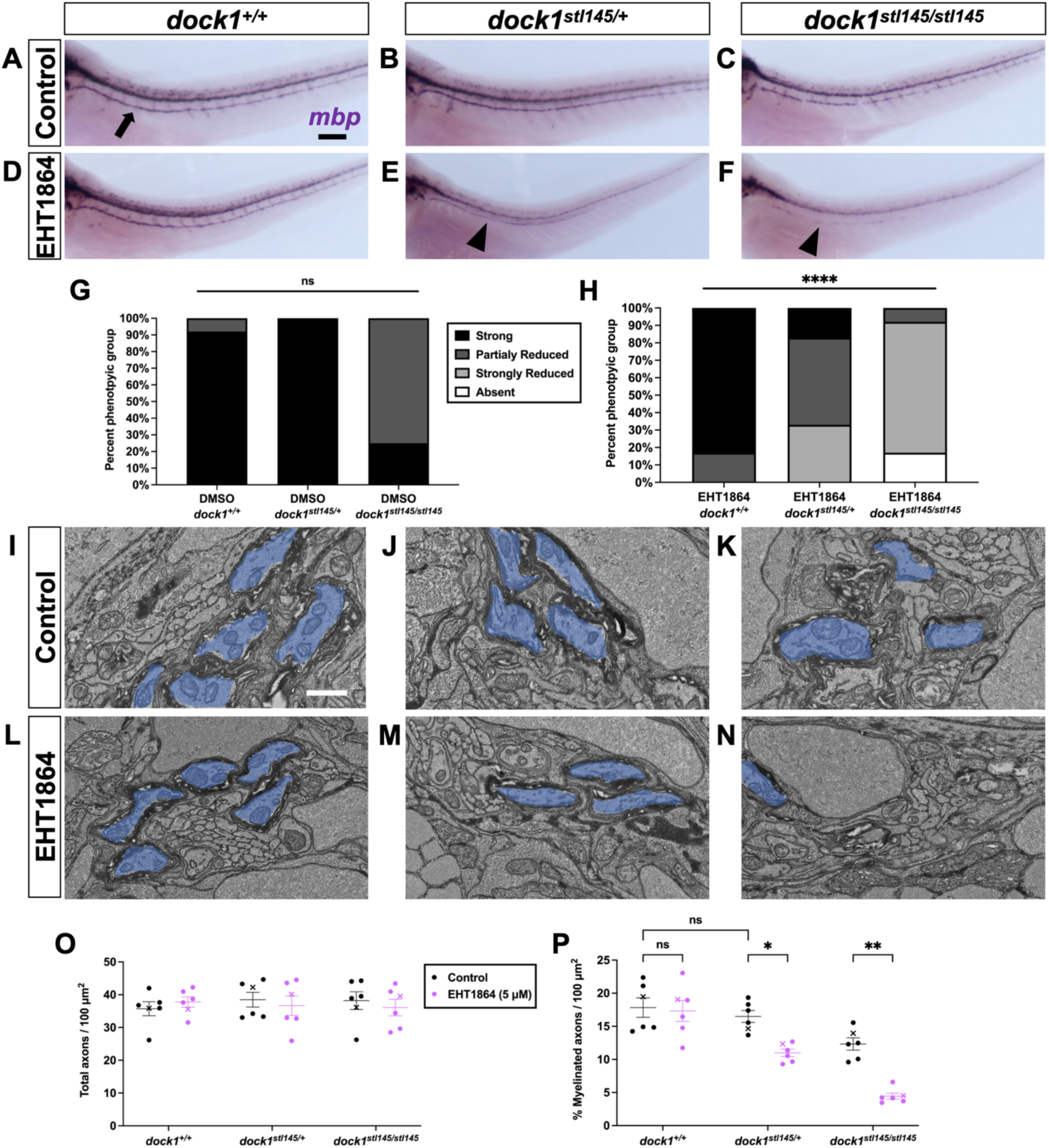
dock1 mutant zebrafish are sensitized to Rac1 inhibition. **(A-F)** Lateral views of larvae showing *mbp* expression by WISH in DMSO control-treated and EHT1864-treated WT *dock1^+/+^*, heterozygous *dock1^stl145/+^*, and homozygous *dock1^stl145/stl145^*mutant zebrafish. **(G, H)** Quantification of *mbp* as seen by WISH in 4 dpf DMSO and EHT1864 zebrafish, compared between phenotypic scores and genotypes. **(I-N)** TEM micrographs of cross-sections of the posterior lateral line, showing myelinated axons pseudocolored in blue, in DMSO control-treated and EHT1864-treated WT *dock1^+/+^*, heterozygous *dock1^stl145/+^*, and homozygous *dock1^stl145/stl145^* mutant zebrafish. **(O, P)** Quantifications of the total axons and myelinated axons per 100 µm^2^ in the posterior lateral line. *n =* 6 fish per genotype (DMSO control) and 6 fish per genotype (EHT1864). **(A-F)** Scale bar = 100 µm, **(I-N)** Scale bar = 1 µm. **(G, H)** Chi-squared analysis. **(O, P)** Two-way ANOVA with Sidak’s multiple comparisons test. *, P < 0.05; **, P < 0.01; ns, not significant.

## Discussion

We previously established Dock1 as an important regulator of developmental myelination in zebrafish and showed that a global mutation in *dock1* results in early developmental hypomyelination (Cunningham et al., 2018). In the present study, we used zebrafish and mouse models to more fully define the role of Dock1 in PNS myelination. In zebrafish, we analyzed adult animals to look beyond development, where we found that Dock1 is vital for the long-term maintenance of myelin health and in repair and remyelination following nerve injury. In a complementary series of experiments in mice, we found that the observations made in global *dock1* zebrafish mutants stemmed from an evolutionarily conserved function of Dock1, where we showed that it acts cell-autonomously in SCs.

### A unique model to study myelin in the adult zebrafish PNS

To look beyond development in zebrafish, we turned to a system that has been characterized but not previously used as an experimental tool, the ZBM. We found that the myelin of *dock1^stl145/stl145^* mutants was indistinguishable from WT in early adulthood but that these mutants had accumulated a significant number of myelin abnormalities at 1 year of age. Next, we used the regenerative capabilities of the ZMB to assess if Dock1 functions in remyelination following nerve injury. We found that mutants regenerated the same total number of axons; however, there was a significant reduction in remyelinated axons 28 days post-transection. These findings expanded our understanding of Dock1’s importance in the zebrafish PNS; however, we could not assign specific functions Dock1 might have in a particular cell type since the experiments were performed in global mutants.

### Dock1 functions cell-autonomously in Schwann cells to regulate PNS myelination

To determine whether Dock1 functions cell-autonomously in SCs and if our findings in zebrafish were evolutionarily conserved in mammals, we generated SC-specific *Dock1* knockout mice using validated and well-characterized mouse lines (Jaegle et al., 2003; Laurin et al., 2008; Leone et al., 2003). Our developmental SC-specific *Dock1* knockout mice had reductions in myelin thickness at P3 during early development and abnormal SC morphology. SC-specific *Rac1* mutant mice have similar phenotypes as SC-specific *Dock1* mutants, including signs of delayed radial sorting and early developmental hypomyelination (Benninger et al., 2007; Guo et al., 2012; Nodari et al., 2007).

Additionally, *Dock1* mutant SCs phenocopy *Rac1* (Benninger et al., 2007; Guo et al., 2012; Nodari et al., 2007) and *Gpr126/Adgrg6* (Mogha et al., 2013) mutant SCs in terms of aberrant cytoplasmic protrusions and accompanying basal lamina trails. When we examined mutant mice at P28, we found that the hypomyelination was no longer present; however, when we looked at one year, we saw a significant increase in myelin abnormalities, similar to what we had observed in fish. It is not uncommon for genes important in development to also play a role in myelin maintenance (Bremer et al., 2011; Decker et al., 2006; Ackerman et al., 2018). That we observe early and late phenotypes in *Dock1* mutants could suggest that the cytoskeletal abnormalities that give rise to developmental myelin defects resolve in early adulthood, perhaps due to compensation but other Dock family members (more on this below), but become dysregulated again in mature animals.

RhoGTPases, including Rac1, regulate many of the signaling pathways in SCs associated with repair and remyelination, including MAP kinases and c-Jun (Harrisingh et al., 2004; Park and Feltri, 2011; Syed et al., 2010). To assess the function of Dock1 in mammalian PNS repair, we performed sciatic nerve transections, a method often used to examine debris clearance and remyelination, which more closely aligns with the ZBM transection model than a nerve crush injury. In *Drosophila*, the ortholog of Dock1, known as CED-5, operates in conjunction with CED-2 and CED-12, homologs of mammalian CrkII and Elmo, respectively. This complex functions as a guanine nucleotide exchange factor (GEF) to activate downstream Rac1 (Ziegenfuss et al., 2012). Disruption of CED-2/CED-12 signaling, or a parallel pathway, led to suppression in the engulfment and degradation of cellular debris. Along these lines, when we performed sciatic nerve transection in mice, we found evidence of altered debris clearance at 14 dpi, where nerves from *Dock1* icKO mice had more foamy macrophages and appeared less cellular than controls. When we looked later to assess remyelination, we found that *Dock1* icKO mice exhibited a significant decrease in remyelinated axons 25 days after transection compared to controls, similar to zebrafish ZMB studies, thus demonstrating a crucial and evolutionarily conserved role for Dock1 in SCs during repair and remyelination.

### What are the signaling partners of Dock1?

The Rho-GTPase Rac1 has been extensively characterized for its role in modulating cellular morphological transformations, primarily by orchestrating cytoskeletal dynamics through actin polymerization. This function has implications in SC development, where differential Rac1 expression regulates the timing of SC migration, radial sorting, and myelination (Benninger et al., 2007; Guo et al., 2012; Nodari et al., 2007). Dock1 is known to exert GEF activity on Rac1 (Benninger et al., 2007; Guo et al., 2012; Nodari et al., 2007); however, the relationship between Dock1 and Rac1 signaling has yet to be examined in the context of myelination. Returning to our zebrafish models, we asked whether *dock1^stl145/+^* heterozygotes, whose *mbp* expression and myelin morphology are indistinguishable from WT, would be sensitized to Rac1 inhibition. This was precisely the case, with low-level Rac1 inhibition leading to a reduction of *mbp* expression, the number of myelinated axons in *dock1^stl145/+^* heterozygotes, and an enhancement of the *dock1^stl145/stl145^* mutant phenotypes, revealing a place for Dock1 as a potential interacting partner of Rac1 in developmental PNS myelination.

It is established that Dock1 binds to the adapter protein Elmo1, an interaction that stabilizes the connection with Rac1 and directs the assembled protein complex to the plasma membrane where it regulates the cytoskeleton (Brugnera et al., 2002; Grimsley et al., 2004; Komander et al., 2008; Lu et al., 2004; Lu and Ravichandran, 2006; Mikdache et al., 2020). Despite the fundamental role of Rac1 in SCs, the precise subcellular site of its activation has yet to be determined. The radial sorting abnormalities in *Rac1* mutants are shared with mutants that influence proteins tied to the basal lamina of the SC, like those found in *laminin* mutants (Chen and Strickland, 2003). This observation might suggest a potential abaxonal positioning for Rac1. Conversely, since the SC’s plasma membrane extensions during radial sorting demand the intertwining of processes into axonal bundles, one might also infer that the localization of the active Rac1 signal could be on the adaxonal side, where the SC directly interacts with the axon. Understanding the specifics of this signaling will provide valuable insight into SC development and enhance our understanding of how SCs sort axons.

Since myelination during development is important for the normal function of the PNS, having multiple GEFs regulate this process and intersect at the same pathway could provide built-in redundancy and a biological advantage, permitting radial sorting and myelin to form even if a single GEF functions abnormally. This may help explain why the early developmental phenotypes we observed in zebrafish and mice resolve in early adulthood. Dock1 might operate with other GEFs, which could be upregulated or act redundantly when it is nonfunctional to ultimately control Rac1 levels. This redundancy may come from other members of the Dock1 family, such as Dock7 or Dock8, which have been shown to have roles in regulating SC migration and development (Miyamoto et al., 2016; Yamauchi et al., 2008; Yamauchi et al., 2011).

Proteins upstream of Dock1 have yet to be well defined. It is known, however, that RhoGEFs can be activated by and function downstream of receptor tyrosine kinases (RTKs), and accordingly, Dock1 has been suggested to function downstream of RTKs in several biological contexts (Duchek et al., 2001; Feng et al., 2012). For example, ErbB2 (HER2) interacts with DOCK1 in breast cancer cells (Laurin et al., 2013). In the context of SCs, the ErbB2/3 heterodimer is the most thoroughly investigated RTK pair, exhibiting critical functions across multiple developmental phases, encompassing migration, radial sorting, and myelination (Monk et al., 2015). The developmental stages regulated by ErbB2/3 in SCs require dramatic cell shape changes and process extension, and as previously noted, Dock1 regulates similar cell shape changes in many biological systems. Additionally, ErbB2, through Rac1 and Cdc42, has been shown *in vitro* to activate Dock7 to regulate SC migration (Yamauchi et al., 2008), positioning, ErbB2/3 as a promising candidate for an upstream signaling partner of Dock1. Alternatively, Dock1 functions downstream of chemokine GPCR signaling in endothelial cell migration (Laurin and Côté, 2014), and the Dock1 adapter protein Elmo1, which has been shown to regulate zebrafish PNS myelination (Mikdache et al., 2020), directly interacts with adhesion G protein-coupled receptors Bai1/Adgrb1 and Bai3/Adgrg3 in myoblast fusion (Hamoud et al., 2014; Hochreiter-Hufford et al., 2013). Interestingly, our SC-specific *Dock1* mutant mice phenocopy the abnormal cytoplasmic protrusions overserved in the SCs of mice with a mutated form of Gpr126/ Adgrg6, another adhesion G protein-coupled receptor (Mogha et al., 2013). In the future, it will be interesting to assess if Gpr126/Adgrg6, which is required for timely radial sorting and essential for SC myelination (Monk et al., 2009), is an upstream activator of Dock1.

In summary, our work combines a series of *in vivo* experimental approaches from zebrafish and mice to demonstrate that Dock1 plays an evolutionarily conserved, cell-autonomous function in SCs, and interacts with Rac1 to regulate PNS myelin biology. When Dock1 is not functional in SCs, myelination is dysregulated during development, myelin abnormalities arise in late adulthood, and SCs lose their ability to repair and remyelinate the PNS after nerve injury. These findings provide crucial insights into our understanding of SC and PNS myelin and offer valuable directions for future studies, which will ultimately help us develop better therapeutic interventions.

## Acknowledgments

*Dock1^fl/fl^* mutant mice were a kind gift from Jean-François Côté. We thank Emma Brennan and Adriana Reyes for their assistance with mouse work. We thank Austin Forbes and Tia Perry for caring for our zebrafish and maintaining our facility. We thank Peter Arthur-Farraj for helpful advice related to the nerve injury studies as well as Monk lab members for feedback on the manuscript.

This work was supported by R01NS120651 to K.R.M and T32NS007446 to R.A.D.

Author contributions: Conceptualization, R.A.D and K.R.M; Formal Analysis, R.A.D; Funding Acquisition, R.A.D and K.R.M; Investigation, R.A.D; Methodology, R.A.D and K.R.M; Supervision, K.R.M; Writing - Original Draft, R.A.D; Writing - Review & Editing, R.A.D and K.R.M.

## Materials & Methods

### Zebrafish lines and rearing conditions

All animal experiments and procedures performed for this manuscript were done so in compliance with the institutional ethical regulations for animal testing and research at Oregon Health & Science University (OHSU). *dock1* transgenic zebrafish (Cunningham et al., 2018) are maintained as heterozygotes (*dock1^stl145/+^*), an incross of which yields wild-type, heterozygous, and homozygous viable zebrafish. Zebrafish larvae are fed a diet of rotifers and dry food (Gemma 75) from 5 days post fertilization (dpf) until 21 dpf. From 21 dpf until 3 months, fish are fed using rotifers and dry food (Gemma 150). Adult fish are maintained and fed with brine shrimp and dry food (Gemma 300). For larval zebrafish studies, sex cannot be considered as a biological variable as sex has not yet been determined in this species. For experiments using adult zebrafish, equal numbers of males and females were examined.

### Mouse strains and maintenance

All mice used, *Dock1^fl/fl^* (Laurin et al., 2008), *Dhh^Cre^*(Jaegle et al., 2003) and *PLP^Cre-ERT2^* (Leone et al., 2003) are previously described and validated lines. For experiments using *Dhh^Cre^* (cKO), *Dhh^Cre+^;Dock1^fl^*^/+^ mice were crossed to Dock1*^fl/fl^* mice to generate *Dhh^Cre+^;Dock1^fl/fl^*mice and their sibling controls. For experiments using *Plp^Cre-ERT2^* (icKO), *Plp^Cre-ERT2+^;Dock1^fl^*^/+^ mice were crossed to Dock1*^fl/fl^* mice to generate *Plp^Cre-ERT2+^;Dock1^fl/fl^*mice and their sibling controls. To induce Cre recombination and Dock1 deletion in icKO mice, 2-month-old *Plp^CreERT2-^;Dock1^fl/fl^*(control), and *Plp^CreERT2+^;Dock1^fl/fl^* (icKO) mice were injected for 5 consecutive days with 2 mg/ml of tamoxifen. For all mouse experiments, mice of both sexes were analyzed, and mutants were always compared with littermate sibling controls.

### Genotyping

Zebrafish - *stl145* primers were used to amplify a region of interest by PCR: F: 5’-CATAGGCGTTCTTCACTGAG -3′ and R: 5’-GACAACAGCTGCCTAATCCG -3’. After PCR, a restriction enzyme digest assay was performed, and the resulting fragments were analyzed on a 3% agarose gel. The *stl145* C-to-T mutation disrupts a BstNI site so that the wild-type PCR product is cleaved into 48 and 353 base pair (bp) products, and the mutant PCR product is 401 bp. Mice - The following primers to detect the presence of the alleles: *Dock1^fl/fl^*, 5′-TCAGCAGGCCCAGTTCCTACT-3′; 5′-GCAGAGCTAGGAGTTCATCGTAGTTC-3′, *Dhh^Cre^*, 5’-CCTTCTCTATCTGCGGTGCT-3’; 5’-ACGGACAGAAGCATTTTCCA-3’, *PLP^Cre-ERT2^*, 5’-CACTCTGTGCTTGGTAACATGG-3’; 5’-TCGGATCCGCCGCATAAC-3’. After PCR, the resulting products were analyzed on a 3% agarose gel.

### Zebrafish maxillary barbel transection

Adult zebrafish were anesthetized with 0.16 mg/ml Tricaine diluted in system water, placed onto a SYLGARD™ (Dow Chemical) filled plate, and visualized under a stereomicroscope. A pair of fine forceps was used to grab the distal most tip of the barbel and lift it away from the surface of the fish. The cut was performed by placing a pair of microdissection scissors parallel to the mouth’s surface to ensure consistency in the cut site between animals. Once removed, the barbel was placed into Karnovsky’s fix (2% glutaraldehyde, 4% PFA in 0.1M sodium cacodylate, pH 7.4), kept on ice, and processed as described below. Fish were returned to individually housed tanks to track them during the regeneration period, and the same procedure was repeated 28 days later, this time the maxillary barbels from each side.

### Sciatic nerve transection

Mice were anesthetized with isofluorane before and during surgery. Fur was removed with an electric razor and the sciatic nerve of the right hindlimb was exposed by making a small cut in the skin. The exposed sciatic nerve was quickly crushed with forceps coated in powdered carbon to mark the injury site and then transected at that location. After transection, surgical wounds were sutured with nylon thread and sealed with metal clips. Mice were monitored daily and administered pain-reducing chow (Bio-Serv) during recovery until they were euthanized for experimental endpoints.

### Transmission electron microscopy

Zebrafish - zebrafish larvae and adult barbels were processed as follows. For larvae, zebrafish were anesthetized with Tricaine and then cut between body segments 5 and 6 to control for variability along the anterior-posterior axis. For ZMBs, the structures were removed by placing a pair of microdissection scissors parallel to the skin to ensure a consistent cut as close to the facial surface. Samples were immersed in Karnovsky’s fix (2% glutaraldehyde, 4% PFA in 0.1M sodium cacodylate, pH 7.4) and microwaved (PELCO BioWave processing - Ted Pella) for at 100 W for 1 min, OFF for 1 min, 100 W for 1 min, and OFF for 1 min, 450 W for 20 s and OFF for 20 s. This was repeated five times, and samples were allowed to fix overnight at 4°C. The following day, samples were rinsed 3 times in 0.1 M sodium cacodylate buffer at room temperature, 10 minutes each rinse. A secondary fixative solution of 2% osmium tetroxide was prepared by combining 2 mL of a stock 0.2M sodium cacodylate + 0.2M imidazole solution (pH 7.5) with 2 mL 4% osmium tetroxide. The 2% osmium tetroxide was added to the samples, and they were microwaved - 100 W for 1 min, OFF for 1 min, 100 W for 1 min, OFF for 1 min, 450 W for 20 s, and OFF for 20 s. This was repeated 5 times, and we allowed them to sit for an additional 2 hours at room temperature. The osmium tetroxide was removed, and the samples were washed 3 times with deionized water, 10 minutes per wash. UranyLess (Electron Microscopy Sciences) was then added to the tubes, and the microwave was run - 450 W for 1 min, OFF for 1 min, and 450 W for 1 min. The samples remained in UranyLess overnight at 4°C. The following day, the UranyLess was removed, and the samples were washed 3 times with deionized water, 10 minutes per wash. A series of ethanol:water (25:75, 50:50, 70:30, 80:20, 95:5, and 100:0) solutions were prepared. Samples were then passed through this graded series of increasing ethanol concentrations, 25% EtOH, 50% EtOH, 70% EtOH, 80% EtOH and 95% EtOH, and were microwaved - 250 W for 45 s followed by incubation at room temperature for 10 min for each concentration. Next, they were changed into a 100% EtOH solution and microwaved at 250 W for 1 min, OFF for 1 minute, and then 250 W for 1 minute; then incubated at room temperature for 10 minutes. This step was repeated with the 100% EtOH 2 more times, for 3x in the 100% EtOH. Next, samples were dehydrated using 100% EM grade acetone and microwaved - 250 W for 1 minute, OFF for 1 minute, and 250 W for 1 minute; and incubated at room temperature for 10 min. This step was repeated with the 100% acetone 2 more times, for 3x in the 100% acetone. Next, a 1:1 solution of Araldite 812:100% acetone was added to the samples and allowed to infiltrate at room temperature overnight. The following day, a fresh batch of Araldite 812 was prepared. With the aid of a dissecting microscope, the samples were carefully oriented in molds so that they were properly aligned for sectioning. They sat at room temperature for 4-6 hours in the Araldite 812 before being placed in a 65°C oven and allowed to polymerize for a minimum of 48 hours.

Mice - sciatic nerves were removed from mice and fixed in a modified Karnovsky’s fix (2% glutaraldehyde, 4% PFA in 0.1M sodium cacodylate, pH 7.4) at 4°C overnight. Nerves were pinned down in a SYLGARD filled dished using 0.20 mm insect pins (Austerlitz) to ensure that they fixed straight. 4-0 Nylon sutures were tied around the distal end of the nerve and removed at the time of embedding to ensure correct cutting orientation. Following fixation, nerves were rinsed 3 times, 15 minutes each, in 0.1M Sodium Cacodylate Buffer and then postfixed with 2% Osmium Tetroxide (as described above) overnight at 4°C. Nerves were then dehydrated in a graded ethanol series (25%, 50%, 70%, 95%, 100%) 3x for 20 minutes per solution. An additional 20-minute 50:50 ethanol: propylene oxide and 2x 20-minute 100% propylene oxide dehydrations were performed before overnight incubation in 50:50 Araldite 812:propylene oxide. For 2 days, nerves were switched to a 70:30 and 90:10 Araldite 812:propylene oxide mix and left overnight at 4°C. On the final day, nerves were put in 100% Araldite 812, allowed to sit at room temperature for several hours to allow infiltration, placed in labeled molds, and baked for a minimum of 48 hours at 65°C. For all zebrafish and mouse samples, semithin sections (400 nm) were stained with toluidine blue and viewed on a light microscope (Zeiss AxioImager M2) to ensure quality before cutting for TEM. Ultrathin sections (60 nm) were cut and counter stained with UranyLess (Electron Microscopy Science), and 3% Lead Citrate and then images were acquired on an FEI Tecnai T12 TEM microscope using an Advanced Microscopy Techniques (AMT) CCD camera.

### Whole mount *in situ* hybridization

Zebrafish were fixed in 4% PFA (made in 1X PBS) overnight at room temperature (RT) for 2 hours with shaking. The PFA was replaced with 100% MeOH, 5 x 5 minutes each in 100% MeOH. After the final wash, embryos were stored in 100% MeOH at -20°C until they were ready to be processed. On day 1 of processing, embryos were rehydrated into PBSTw, (50% PBSTw – 70% PBSTw – 100% PBSTw) with 5-minute washes each, followed by 4 x 5-minute PBSTw washes at RT with shaking. Samples were placed in 1:2000 ProtK liquid stock (20 mg/ml) in PBS without shaking for 55 minutes. The ProtK was removed, followed by 2x 5-minute PBSTw washes to remove the ProtK. Samples were postfixed in 4% PFA for 20 min at RT with shaking. The PFA was removed and there were 5 x 5-minute washes in PBSTw at RT with shaking. Next, the samples were prehybridized in 400 µL Hyb(+) solution for 1-2 hours at 65°C. Tubes were kept on their sides to ensure adequate exposure to the solution. Next, 400 µl of probe diluted in Hyb(+) was added to each tube and left overnight at 65°C, with the tubes on their sides. On day 2, all the solutions used were preheated to 65°C before adding them to the samples, and all washes were done at 65°C, taking care to ensure the samples were not allowed to cool down. The probe was removed and saved, 100% Hyb was added, and the samples were left to sit for 5-10 minutes. Next, a series of liquid changes were performed: 75% Hyb:25% 2X SSCTw 5 min at 65°C, 50% Hyb:50% 2X SSCTw 5 min at 65°C, 25% Hyb:75% 2X SSCTw 5 min at 65°C, 2X SSCTw 2 x 30 min at 65°C, 0.2X SSCTw 2 x 30 min at 65°C, and MABTr 10 min at RT, shaking tubes on side. A blocking solution was prepared by combining: 2% blocking reagent in MAB + 0.2%Triton +10% sheep serum. The block was added to the samples and incubated for 1-2 hours at RT, while shaking tubes on their side. Next, the block was removed and replaced by Anti-Dig AP Fab fragments, diluted 1:2000 in blocking solution and left overnight at 4°C with shaking. On day 3, the Anti-Dig AP Fab fragment solution was removed and MABTr washes were performed: 6x 30 minutes each at RT with shaking. The MABTr was removed, and AP/NTMT buffer was added and allowed to sit for 10 minutes at RT with shaking. The samples were then incubated in a solution of: AP buffer + NBT (2.2 µl/ml AP buffer) + BCIP (1.6 µl/ml AP buffer) and covered in foil as the reaction is light sensitive. The reaction proceeded for 2-3 hours until the lateral line was visible under a light microscope, and the reaction was stopped by doing 3 quick washes in PBSTw. The samples were then postfixed in 4% PFA for 30 min at RT. The PFA was removed, and samples were passed through a 30% - 50% - 70% glycerol series, moving on to the next one after they sank to the bottom. Samples were stored in 70% glycerol at 4°C until they were ready to be imaged. Samples were mounted onto slides, suspended in 70% glycerol, and brightfield imaged using a Zeiss Discovery.V8 stereomicroscope.

### EHT1864 treatment

Zebrafish larvae were treated with 0.003% phenylthiourea (PTU) at 24 hours post fertilization (hpf) to inhibit pigmentation. At 48 hpf, immediately before treatment, zebrafish were manually dechorionated with a pair of #5 forceps. Zebrafish were treated with DMSO or 5μM EHT1864 (3872, Tocris), in embryo media with 0.003% PTU from 48-98 hpf. At 96 hpf, larvae were anesthetized with Tricaine and processed for *in situ* hybridization or TEM as described.

### Western blotting

Mice were euthanized, and sciatic nerves from both legs were harvested, placed together in labeled 1.7 mL tubes, and immediately flash-frozen on dry ice and stored at - 80°C until the protein extraction occurred. The sciatic nerves were thawed, and a solution of RIPA buffer (50mM Tris HCl, pH 8.0, 150mM NaCl, 1% NP-40, 0.5% Sodium deoxycholate, 0.1% SDS, 1mM EDTA, 0.5mM EGTA) containing protease inhibitor (11836153001, Roche) was added to the nerves. A sterile tissue homogenizer was attached to a Ryobi drill press and the nerves were homogenized by moving the homogenizer up and down in the dounce 30 times until the nerve appeared completely homogenized. The dounce was kept in a beaker of ice water during this process and care was taken to ensure that the tissue remained cold throughout homogenization. The samples were allowed to rest on ice for 10 minutes, then spun for 15 minutes at 15,000 rpm in a 4°C cooled centrifuge. The supernatant was then removed and added to a new tube. A Bradford protein assay was performed to ensure equal protein concentrations in our samples before proceeding. BCA standards are combined with MQH_2_O and 1 mL of Coomassie Plus. The standards used were 750 µg/mL, 500 µg/mL, 250 µg/mL, 125 µg/mL, 65 µg/mL and 0 µg/mL. For each tube, 5 μL of sample and 495 µL of MQH_2_O were added to a cuvette, along with 1 mL of Coomassie plus. Samples were measured on a Nanodrop spectrophotometer following the measurement of a blank sample. The spread between the lowest and highest protein concentrations was < 5%. Next, 1 part Laemmli buffer was combined with 4 parts of protein w/ RIPA buffer and samples were thoroughly mixed. Samples were heated for 5 minutes to 95°C and briefly spun. The gel tank was assembled and a 4-12% Bis-Tris Gel (NP0335BOX, Invitrogen) was loaded with ladder 9 and 25μL of sample per lane. The gel was run at 150V for 1 hour. The gel was then removed and placed into a sandwich with a PVDF membrane (IPVH00010, Thermo Fisher Scientific), and sponge pads in a gel blotting cassette (A25977, Thermo Fisher Scientific). The transfer was run at 20V for 1 hour. The membrane was placed in a black box and washed with 1x TBS with 0.1% Tween-20 (TBST) for 10 minutes. The membrane was then blocked with 5% milk powder in 1x TBST on a shaker @ RT for 1 hour. The membrane was then transferred into Dock1 primary antibody (1:1000, 23421- 1-AP, Proteintech) made in 1x TBST with 2% BSA and incubated overnight at 4°C with shaking. The following day, the primary antibody was removed and saved. The membrane was washed 3x, 5 minutes each, with TBST and following the final wash, HRP conjugate goat anti-rabbit secondary (7074, Cell Signaling), 1:2000 in 1x TBST with 2% milk powder, was added. The membrane was incubated at room temperature for 2 hours and rinsed 3x with TBST and 1x with TBS. The membrane was visualized using a chemiluminescence reaction (34080, Thermo Fisher Scientific) and imaged with a Syngene GBox iChemiXT. Following imaging, membranes were washed with TBST and re-probed with HRP conjugated β-actin (A3854, Sigma-Aldrich). Densitometric analysis was performed in Fiji by quantifying the intensity of the Dock1 protein bands relative to the β-actin loading control and then normalized relative to the controls.

### Morphological characterizations

For determining the % of axons with abnormal myelin in zebrafish at 4- and 12-months of age, the number of myelinated axons with disrupted myelin sheaths (splitting, degeneration) was divided by the total number of myelinated axons. For TEM analysis in mice: To calculate g-ratios, we manually measured axon diameter and axon-plus-myelin diameter in ImageJ. We measured a minimum of 100 axons from 3 ∼2000 μm2 regions of each sciatic nerve selected at random. The measurements were taken with the observer blind to treatment. To determine the % of axons with abnormal profiles in mice, the number of abnormally myelinated axons (outfolding, degeneration, decompaction) was divided by the total number of myelinated axons. In addition, the percentage of abnormal Remak bundles and the number of degenerating axons compared to controls were included. For quantification of *in situ* hybridization, larval zebrafish were blinded, imaged, and assigned values of “strong,” “partially reduced,” “strongly reduced,” and “none” based on *mbp* expression in the lateral line.

### Statistical analysis

All statistical analyses were performed using GraphPad Prism 10. For zebrafish barbel morphometric analysis, cross sections of the entire barbel were analyzed. For an analysis of the effect of two variables (genotype and age) or (genotype and control/injured), two-way ANOVA was used with Tukey’s or Sidak’s multiple comparisons test to analyze the effect of genotype and experimental condition compared to controls. When comparing multiple experimental groups to the same control group, a one-way ANOVA with a Brown-Forsythe test was used. When comparing one experimental group to a control, we used an unpaired t test with a Welch’s correction. For quantifying *mbp* expression by in situ, an average for each score per genotype and condition was calculated, and a Chi-squared analysis was performed to determine significance. P values shown are represented as follows: *, P < 0.05; **, P < 0.01; ***, P < 0.001; ****, P < 0.0001; ns, not significant.

---

**Supplemental Figure 1.**
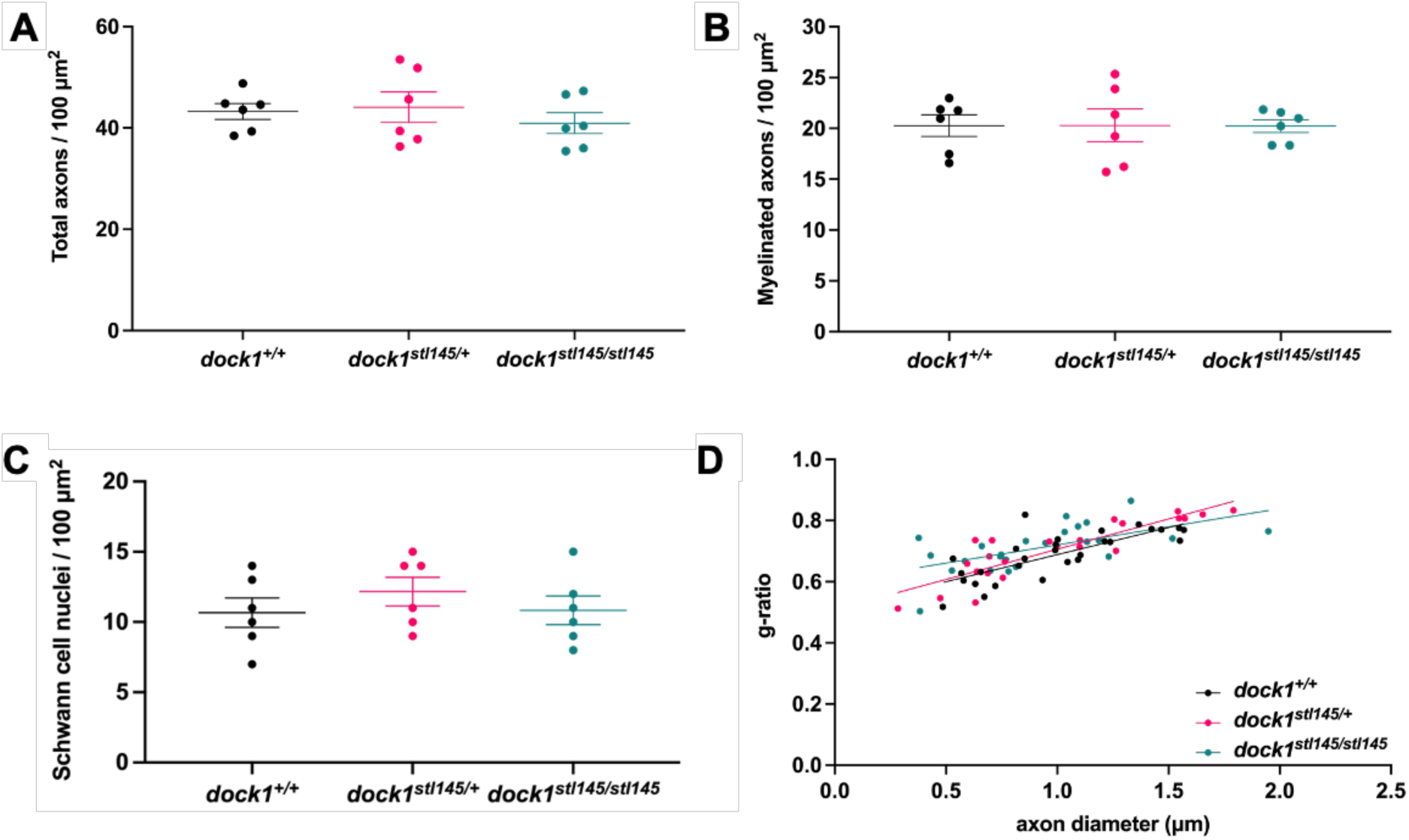
*dock1* mutant zebrafish don’t exhibit myelin defects at 4 months. **(A-D)** Quantifications of the total number of axons, the number of myelinated axons, the number of Schwann cell nuclei, and g-ratio were obtained from analyzing TEM micrographs. None of these analyses revealed significant differences between WT and mutant zebrafish. **(A-C)** One-way ANOVA with Brown-Forsythe test.

**Supplemental Figure 2.**
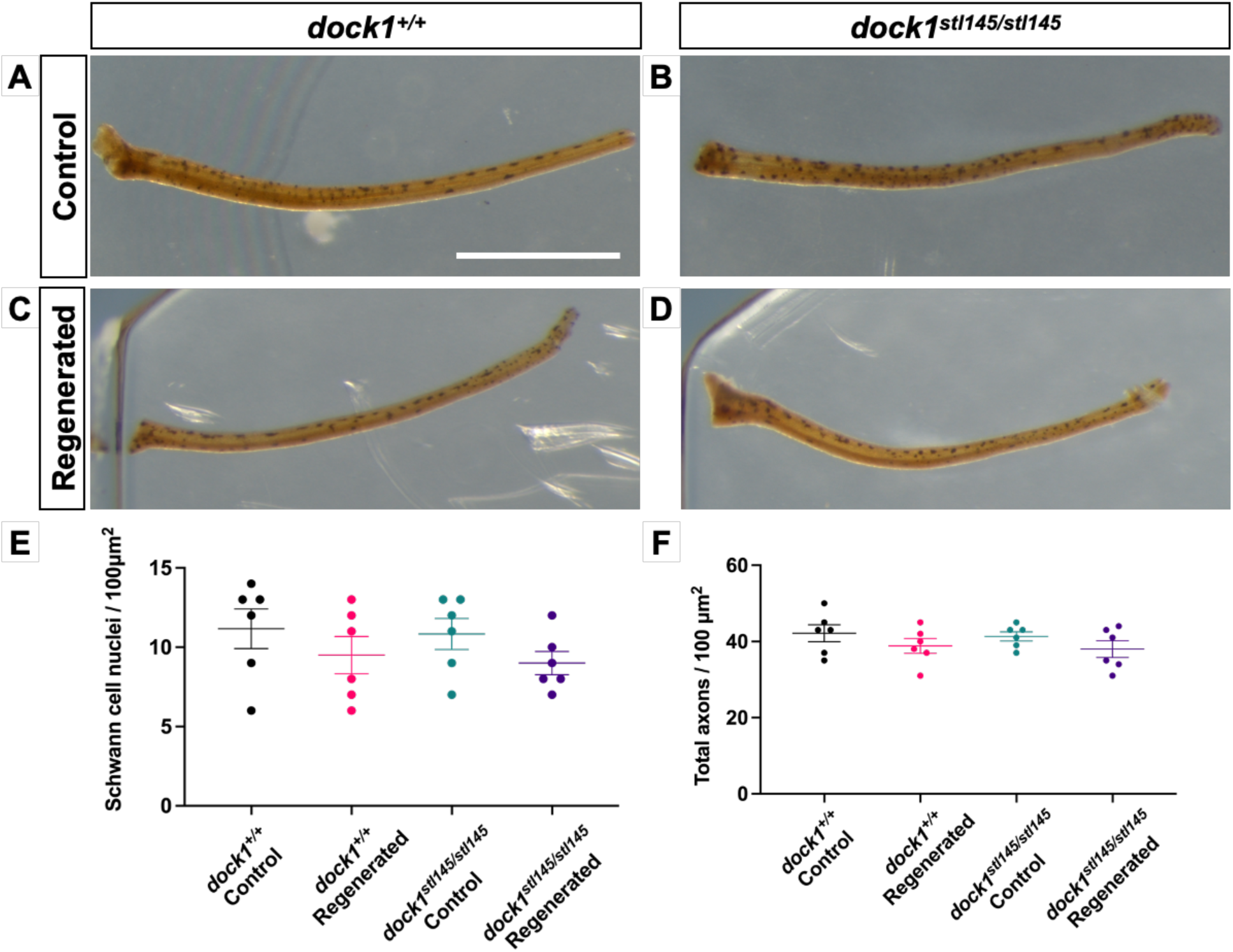
Adult *dock1* mutant zebrafish regenerated barbels are grossly indistinguishable from WT. **(A-D)** Maxillary barbels from uninjured 4-month-old WT and *dock1* mutant zebrafish (A, B) taken at the same time as 28-day regenerated control and *dock1* mutants (C,D) were harvested. **(E, F)** Quantifications from TEM micrographs found no significant differences in the number of SC nuclei or total axon count between genetic and experimental groups. **(A-D)** Scale bar = 1 mm **(E, F)** One-way ANOVA with Brown-Forsythe test.

**Supplemental Figure 3.**
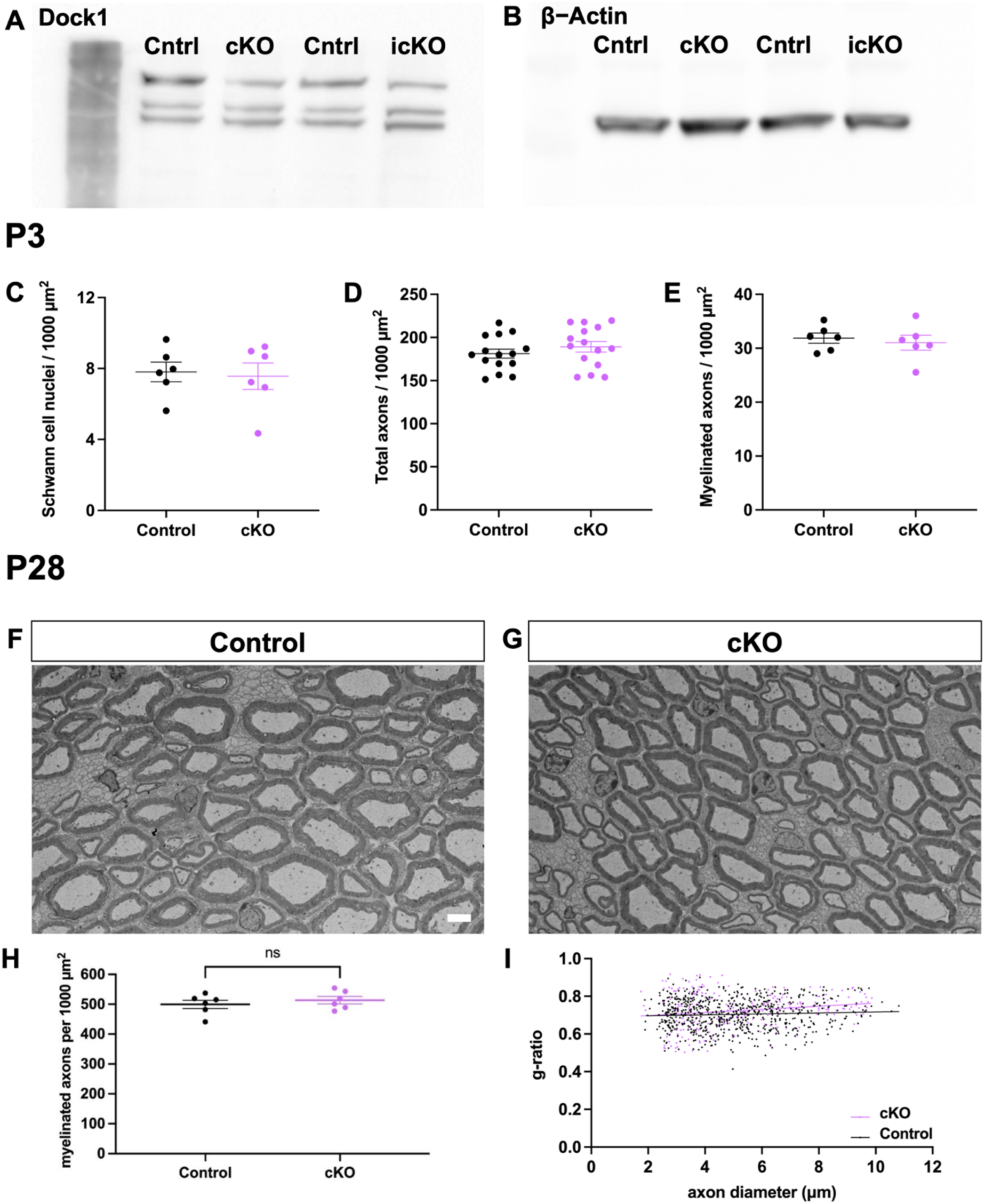
The myelin phenotype observed in *Dock1* mutants at P3 resolves by P28. **(A, B)** Full western blot showing Dock1 **(A)** β-Actin **(B)** protein levels. **(C-E)** Quantifications obtained from analyzing TEM micrographs showing the number of SC nuclei, the total number of axons, and the number of myelinated axons. None of these analyses revealed significant differences between control and cKO mice at P3. **(F, G)** TEM micrographs of control and *Dock1* cKO sciatic nerves at P28. **(H, I)** Quantifications of myelinated axon count and g-ratios obtained from P28 TEM micrographs reveal no significant differences. **(F, G)** Scale bar = 4 µm **(C-E, H)** Unpaired t test with Welch’s correction. ns, not significant.

**Supplemental Figure 4.**
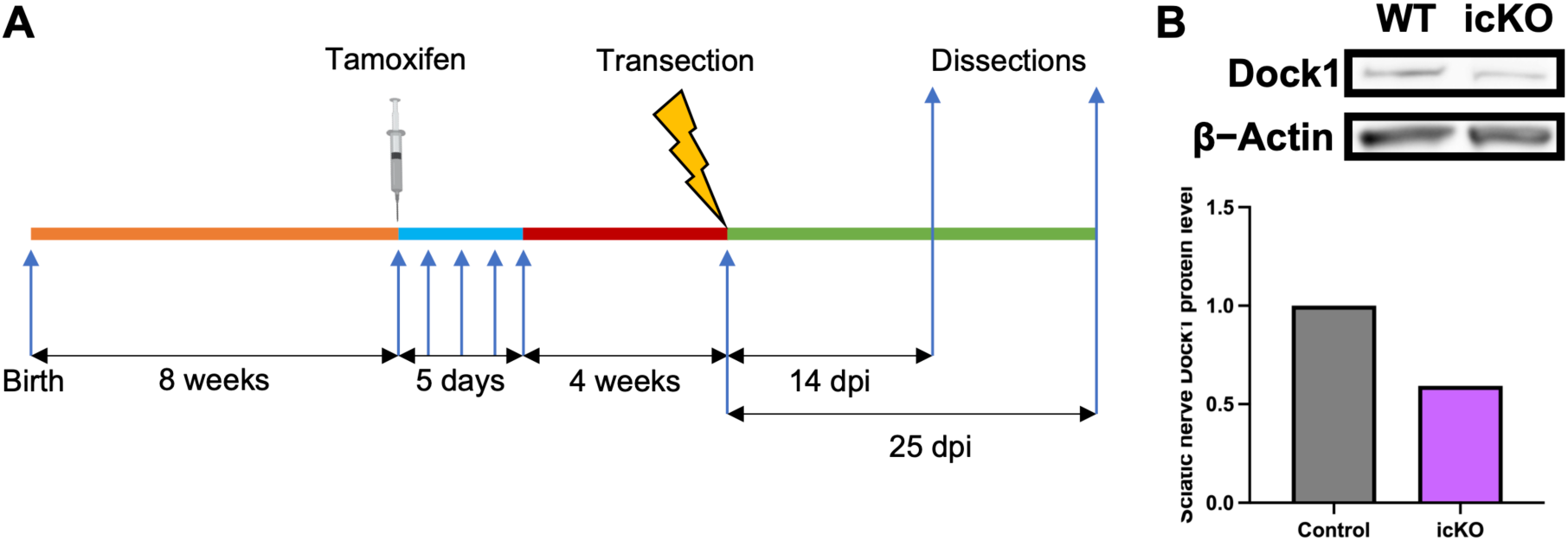
An inducible Schwann cell specific *Dock1* mutant mouse to study SC repair. **(A)** Schematic representation showing the experimental timeline for Sciatic nerve injury studies using the icKO mice. **(B)** Western blot showing sciatic nerve Dock1 and β-actin protein levels from control and *Dock1* icKO animals and quantification of normalized protein levels.

